# Genetically-based adaptive trait shifts at an expanding mangrove range margin

**DOI:** 10.1101/2021.07.17.452777

**Authors:** John Paul Kennedy, Giles N. Johnson, Richard F. Preziosi, Jennifer K. Rowntree

**Affiliations:** Ecology and Environment Research Centre, Department of Natural Sciences, Faculty of Science and Engineering, Manchester Metropolitan University, Manchester, UK; Department of Earth and Environmental Sciences, Faculty of Science and Engineering, University of Manchester, Manchester, UK

**Keywords:** Atlantic Florida, *Avicennia germinans*, climate change, common garden experiment, foundation species, mangrove restoration, range expansion

## Abstract

1. Many species are expanding beyond their distributional range margins in response to a warming planet. Due to marginal environmental conditions and novel selection pressures, range margins may foster unique genetic adaptations that can better enable species to thrive under the extreme climatic conditions at and beyond their current distributional limits. Neotropical black mangrove (*Avicennia germinans*) is expanding poleward into temperate salt marsh along Atlantic Florida, USA, with field evidence of adaptive trait shifts within range-margin *A. germinans* populations. However, whether these adaptive shifts have a genetic basis remains to be answered.
2. We monitored twenty *A. germinans* maternal cohorts from areas in both the Atlantic Florida range core and margin in a greenhouse common garden with annual temperatures analogous to range-margin conditions. We measured variation in a series of phenotypic traits starting at initial planting of field-collected propagules and continuing until two years development.
3. Maternal cohorts from the Atlantic Florida range margin consistently outperformed those from the range core throughout the experiment. Range-margin cohorts survived in greater numbers, established faster, and were less stressed under winter chilling and sub-zero temperatures that are often reached at the Atlantic range margin, but not within the range core. Range-margin cohorts did not grow taller, but instead invested more into lateral growth and biomass accumulation that presumably reflects adaptation to their colder and open canopy environment. Range-margin cohorts also exhibited leaf traits consistent with greater resource acquisition that may compensate for a shorter growing season and reduced light quality at higher latitude.
4. *Synthesis*. We confirmed that there is a genetic basis to adaptive trait shifts towards an expanding mangrove range margin. Our results suggest that genetically-based phenotypic differences better enable these range-margin mangroves to thrive within their stressful environment and may facilitate further poleward expansion in the future. In addition, our documentation of adaptive trait variation among maternal cohorts of an ecologically-important mangrove foundation species, quantitative data that is lacking for mangroves, should help inform mangrove restoration initiatives.

## 1 | INTRODUCTION

Distributional range margins are often defined by a species’ inability to tolerate conditions beyond these boundaries (Brown, 1984). However, in response to a warming planet, these boundaries are expanding poleward for many species (Chen, Hill, Ohlemüller, Roy, & Thomas, 2011; Osland et al., 2021; Pecl et al., 2017), with individuals that inhabit present-day range margins inherently at the forefront of this change. Due to marginal environmental conditions and novel selection pressures, individuals at range margins may exhibit strong genetic divergence and significant phenotypic differences from conspecifics within more benign portions of their range (Chuang & Peterson, 2016; Hardie & Hutchings, 2010). Understanding whether these unique range-margin genotypes are better able to thrive under the extreme climatic conditions at and beyond their current distributional limits can provide important insights into how species may respond to climate change (Nadeau & Urban, 2019; Rehm, Olivas, Stroud, & Feeley, 2015).

Evaluating genetic and phenotypic changes towards expanding range margins of plant foundation species will be particularly informative because of the direct influence of these species on ecosystem structure and function (Ellison et al., 2005). Hence, insights into how foundation species will respond to climate change will inevitably inform predictions about responses of entire ecosystems (Bernhardt & Leslie, 2013). A well-documented example of foundation species undergoing climate-driven range expansion is that of mangroves at their poleward range margins (Armitage, Highfield, Brody, & Louchouarn, 2015; Cavanaugh et al., 2014; Cohen et al., 2020; Fazlioglu, Wan, & Chen, 2020; Osland, Day, et al., 2017; Saintilan, Wilson, Rogers, Rajkaran, & Krauss, 2014; Whitt et al., 2020).

Mangroves are (sub)tropical, intertidal woody plants of significant ecological importance to coastal ecosystems (Lee et al., 2014) and a central component to a growing number of coastal rehabilitation and restoration initiatives (Friess et al., 2019; Waltham et al., 2020). Their distributional limits are defined by region-specific climatic thresholds in minimum temperatures and/or precipitation (Duke, Ball, & Ellison, 1998; Osland, Feher, et al., 2017). Along Atlantic Florida, USA, the northern extent of mangroves is controlled by a gradient in minimum winter temperatures that drives a transition from the southern range core of dense, closed canopy mangrove forests to the northern range margin of sparsely-populated mangrove patches within a landscape of temperate salt marsh (Cavanaugh et al., 2018; Osland, Feher, et al., 2017). Milder winters for several decades are linked to ongoing mangrove proliferation at this range margin (Cavanaugh et al., 2014; Rodriguez, Feller, & Cavanaugh, 2016) and further poleward expansion is forecast as freeze events become less common (Cavanaugh et al., 2019, 2015).

Neotropical black mangrove, *Avicennia germinans*, is the predominant mangrove species at the Atlantic Florida range margin (Lonard, Judd, Summy, DeYoe, & Stalter, 2017). Range-margin populations of *A. germinans* exhibit clear genetic differences from those directly south within the continuous range core (Kennedy, Preziosi, Rowntree, & Feller, 2020) and are the predominant source of new recruits to northern areas beyond this species’ present-day distribution (Kennedy, Dangremond, et al., 2020). Field samples from these range-margin *A. germinans* also demonstrate shifts towards phenotypic traits consistent with greater cold tolerance compared to range-core conspecifics (Cook-Patton, Lehmann, & Parker, 2015; Kennedy, Preziosi, et al., 2020), with similar shifts observed at *A. germinans* range margins in the Gulf of Mexico (Madrid, Armitage, & López-Portillo, 2014; Méndez-Alonzo, López-Portillo, & Rivera-Monroy, 2008). Yet, we lack an understanding of whether these phenotypic differences in range-margin *A. germinans* have a genetic basis or are plastic responses to their marginal environmental conditions. Extensive trait plasticity in response to environmental variation is well documented in mangroves (e.g., Feller et al., 2010; Lovelock, 2008; Vovides et al., 2014), while relatively few studies provide evidence for genetically-based adaptive trait variation.

Common garden experiments are a tool to address this knowledge gap as their uniform environment allows for tests of genetic effects while controlling for trait plasticity (Hoffmann & Sgró, 2011). Furthermore, common gardens with environmental conditions analogous to those that restrict a species’ distribution can provide additional insights into how genetically-based trait variation better suited to tolerate these conditions varies geographically within a species (Alberto et al., 2013; Warwell & Shaw, 2017). In this study, we monitored *A. germinans* maternal cohorts from areas in the Atlantic Florida range core and margin in a greenhouse common garden with annual temperatures that resembled those at the Atlantic Florida range margin. We assessed differences in a series of phenotypic traits starting at initial planting of field-collected propagules and continuing until two years development.

Our aim was to determine whether there is a genetic basis to previous field observations of adaptive trait shifts in *A. germinans* towards its expanding Atlantic Florida range margin (as outlined above). We predicted that, compared to range-core cohorts, (1) field-collected propagules from range-margin cohorts would survive in greater numbers and establish faster. (2) Range-margin cohorts would be less stressed under winter temperatures, which would result in (3) greater growth and biomass accumulation over the two-year experiment. (4) Range-margin cohorts would exhibit more conservative leaf traits (i.e., smaller, increased dry-matter content, reduced specific leaf area) to better tolerate marginal temperature conditions. Our documentation of adaptive trait variation among maternal cohorts of an ecologically-important mangrove species should provide not only insights into dynamics at expanding range margins, but also help inform mangrove restoration initiatives.

## 2 | MATERIALS AND METHODS

### 2.1 | Field sampling

We focused our sampling at the lowest level of genetic inheritance for our studied species (i.e., maternal cohorts). *Avicennia germinans* is a hermaphroditic, insect-pollinated tree or shrub that produces cryptoviviparous propagules (Lonard et al., 2017). Along Atlantic Florida, *A. germinans* exhibit a mixed-mating system with relatively high rates of self-fertilisation (Kennedy, Sammy, Rowntree, & Preziosi, 2021). As such, the maternal cohorts monitored in this research are a mixture of both selfed and outcrossed progeny, with outcrossed progeny being either full- or half-siblings.

On 7 – 8 October 2017, we collected mature *A. germinans* propagules systematically from around the entire canopy of maternal trees located in both the Atlantic Florida range core, where mangroves are the dominant coastal foundation species, and the range margin, where salt marsh vegetation is dominant (Figure 1A). We collected from three range-core and three range-margin sites, across similar geographic expanses (inter-site distances: 47.6 – 97.1 km for range core; 33.2 – 71.4 km for range margin), to include a broader representation of genetic variation across these areas (Figure 1A). Annual minimum temperatures decline with latitude across our sampling area, with temperatures <10°C, a threshold shown to induce chill stress in *A. germinans* seedlings (Devaney, Pullen, Feller, & Parker, 2021), common only at the range-margin sites (Figure 1B). All propagules collected from each maternal tree were stored together in one labelled plastic bag during field collections and then transported to the greenhouse facility at Manchester Metropolitan University in Manchester, UK (53.4713°N, 2.2412°W). Propagules remained dry and intact during transport and planting began 10 days after collection (see *2.3 Common Garden Experiment*).

**FIGURE 1.**
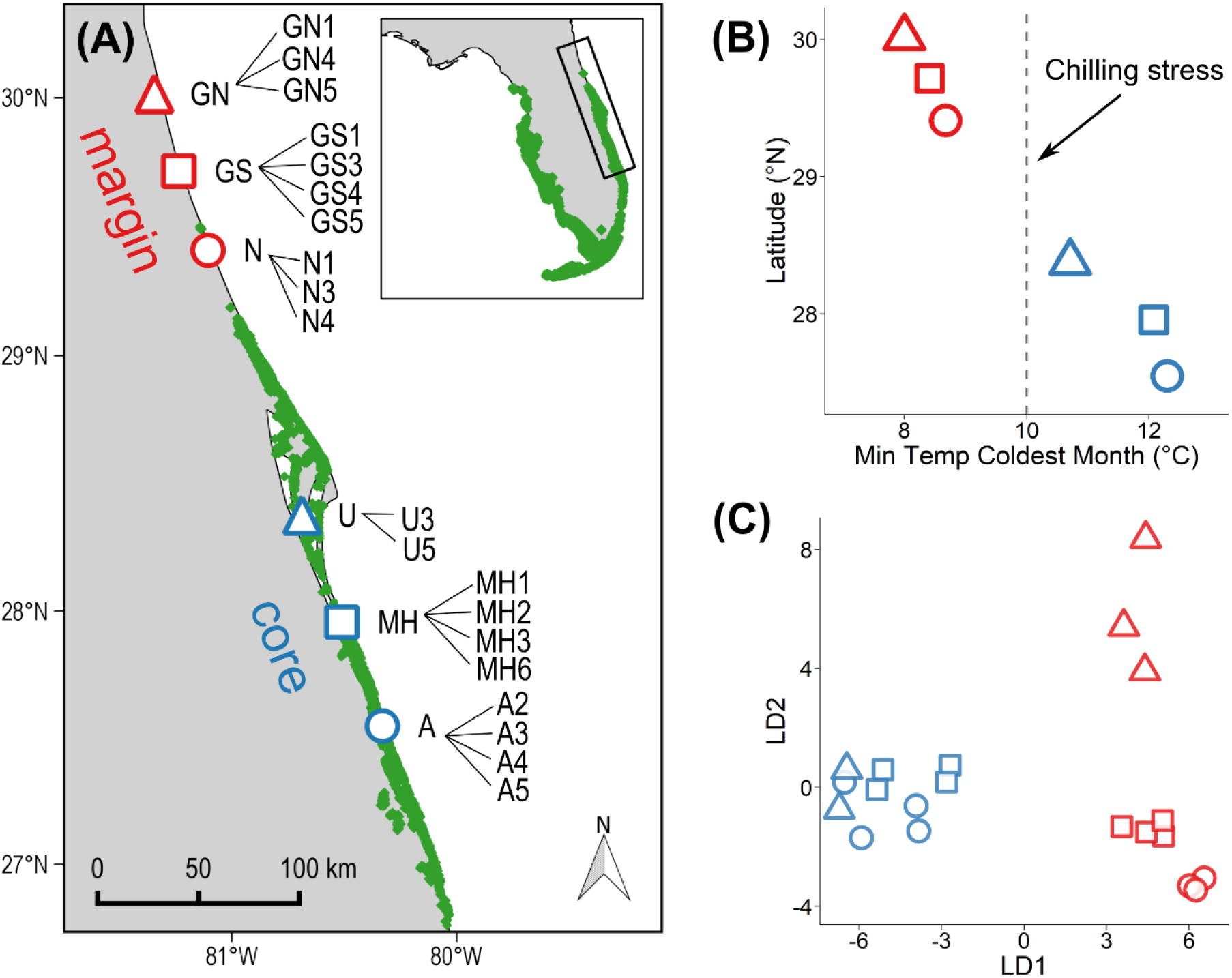
Field collections of *Avicennia germinans* propagules from Atlantic Florida, USA, for a greenhouse common garden experiment at Manchester Metropolitan University, Manchester, UK. (A) Twenty maternal cohorts from six collection sites were included in the experiment (n = 10 from range core, n = 10 from range margin). Mangrove distribution shown in green (Giri et al., 2011) (B) Latitudinal decline in annual minimum temperatures (1970–2000) across the sampled sites, with chilling stress (<10°C) common only at range-margin sites. Temperature data from WorldClim2 (Fick & Hijmans, 2017). (C) Discriminant analysis of principal components (DAPC) of the 20 maternal tree genotypes that demonstrates a clear separation between range core and margin. Throughout the figure, blue shapes depict range-core sites and cohorts, and red shapes depict range-margin sites and cohorts.

### 2.2 | Maternal tree genotyping

During field sampling, we also collected a leaf from each maternal tree to generate their multi-locus genotypes with 12 nuclear microsatellite loci as outlined elsewhere (Kennedy, Preziosi, et al., 2020). We visualised genetic differences among maternal trees with a discriminant analysis of principal components (DAPC) (Jombart, Devillard, & Balloux, 2010) in the adegenet 2.1.1 R-package (Jombart & Ahmed, 2011). For this analysis, we retained nine principal components, which explained ~90% of the total variance, identified two clusters, and retained all five discriminant functions.

### 2.3 | Common garden experiment

Our planting trays permitted the inclusion of 20 maternal cohorts in the common garden experiment, with range-core (n = 10) and range-margin (n = 10) cohorts equally represented (n = 2 – 4 cohorts per collection site) (Figure 1A). The experiment consisted of two components: (1) an establishment phase that monitored propagule development into seedlings until eight months post-planting (20 maternal cohorts x 30 biological replicates = 600 total propagules), and (2) a subsequent growth phase that monitored a random subset of these seedlings until two years post-planting (20 maternal cohorts x 12 biological replicates = 240 total seedlings). We used a randomised complete block design for each component, with one offspring from each of the 20 maternal cohorts present within each block (i.e., replicate planting tray). Greenhouse temperature and humidity were continuously monitored at 30-minute intervals with iButton data loggers (Measurement Systems Ltd, Newbury, UK). We set greenhouse temperatures to resemble those at the Atlantic Florida range margin based on mean monthly values (1981 – 2010) from St Augustine Lighthouse, Florida (29.8°N), obtained from the National Centers for Environmental Information (https://www.ncdc.noaa.gov/cdo-web/datatools). In addition, we superimposed supplemental grow lights (54,800 lm, PLANTASTAR 400 W E40; OSRAM, Munich, Germany) onto natural light throughout the day. The duration of this supplemental light was set each month to match mean monthly day length, also at St Augustine Lighthouse, based on data from the Earth System Research Laboratories (https://www.esrl.noaa.gov/gmd/grad/solcalc/).

Due to the location of the greenhouse facility in Manchester, UK (53.4°N), summer months did experience greater photoperiods than those at the Atlantic Florida range margin (e.g., in June, greenhouse plants experienced ~3 hours more sunlight).

On 18 October 2017, 10 days after collection, we began floating field-collected propagules in a saline water solution (~15 ‰ Instant Ocean® Sea Salt) for one week, an optimal duration for seedling productivity (Simpson, Osborne, & Feller, 2017). On 25 October, we towel dried propagules and measured three size metrics to account for variation in maternal investment, specifically weight (g), length (mm), and width (mm). All three metrics exhibited strong positive correlations (Pearson’s correlation, r = .76–.88, p < .001), so we decided to use propagule weight as our measure of maternal investment. Propagules were then planted in 7 × 7 × 6.5 cm square pots (LBS Horticulture, Colne, UK) filled with a 3:1 mixture of low nutrient commercial potting soil (Levington F1 Seed and Modular Compost; LBS Horticulture, Colne, UK) and sharp horticultural sand (RHS Sharp Sand; LBS Horticulture, Colne, UK), with no subsequent nutrient additions, and placed into 30 replicate trays (Gratnells shallow trays, 42.7 × 31.2 × 7.5 cm; YPO, Wakefield, UK). We added a saline water solution (~15 ‰ Instant Ocean® Sea Salt) to 3 cm depth within the trays to maintain soil saturation, and additional fresh water was added each week to return to this volume. Pots were also misted periodically with fresh water to ensure propagules remained hydrated. Every two weeks, trays were systematically rotated around the greenhouse and salinity was measured from six haphazardly-chosen trays with a handheld refractometer (VWR International, Lutterworth, UK). Complete water changes were performed at the end of each month. We determined that propagules had established as seedlings upon appearance of their first true leaves (Finney, 2011). Once the first seedling established, we began monitoring time to establishment for each propagule on a weekly basis until 35 weeks post-planting when 98.5% of surviving propagules had established. We also documented propagule mortality throughout this period.

On 10 July 2017, 8 months post-planting, we measured height (cm) and total growth as height plus length of any lateral shoots (cm) for all surviving seedlings. Then, on 18 July 2017, a random subset of 12 surviving seedlings from each of the 20 maternal cohorts was transferred to larger 11 × 11 × 12 cm square pots (LBS Horticulture, Colne, UK) filled with a fresh mix of 3:1 potting soil and sand (as detailed above), with no subsequent nutrient additions, and placed into 12 replicate trays (Garland square garden tray, 60 × 60 × 7 cm; LBS Horticulture, Colne, UK). A saline water solution (~15 ‰ Instant Ocean® Sea Salt) was added to 4 cm depth within the trays to ensure soils remained moist, additional fresh water was added each week to return to this volume, and plants were misted periodically. Every month, trays were systematically rotated around the greenhouse and salinity was measured from all trays. Complete water changes were performed every two months. We measured height (cm) and total growth (cm) for all plants at 10, 12, 14, 20, and 24 months post-planting, and documented plant mortality throughout this period.

To evaluate plant stress, we measured maximum quantum yield of photosystem II (F_v_/F_m_) with a FluorPen FP 100 (Photon Systems Instruments, Drásov, Czech Republic). F_v_/F_m_ is an indicator of plant photosynthetic performance and a widely-used diagnostic to measure plant tolerance to environmental stress, with optimal values generally between 0.8 and 0.85 (Baker, 2008; Maxwell & Johnson, 2000). Measurements were taken on a single, undamaged leaf from the youngest, fully-mature leaf pair for all surviving plants at three time points during the experiment: (1) at the start of the second winter (Dec 2018; 13 months post-planting), (2) during a series of nights with sub-zero temperatures (Feb 2019; 15 months post-planting), and (3) at the end of the second summer (Sept 2019; 22 months post-planting). Measurements were taken on three separate dates at time point 1 (4, 11, 18 Dec 2018) to establish baseline F_v_/F_m_ values before the onset of colder winter temperatures, and a similar approach was taken at time point 2 (measurements on 1, 2, 3 Feb 2019) to assess plant responses over the course of a cold event. Values were highly correlated among measurement dates in December (r = .83–.88, p < .001) and February (r = .89–.92, p < .001), so we used mean values across these dates for statistical analyses. Measures for time point 3 were taken only on 5 September 2019 to assess plant performance under more benign conditions. All measurements were taken in the evening on plants that had been dark-adapted for at least two hours.

From 30 October to 7 November 2019, 24 months post-planting, plants were harvested and dried to obtain biomass measurements. Plants were cut at soil level, divided into roots, shoots, and leaves (roots were gently washed with fresh water to remove sediment), and then dried at 60°C for three days until constant weight (g). Prior to harvest, we collected one leaf from each surviving plant to measure functional traits. Leaves were undamaged and from the youngest, fully-mature leaf pair. We measured fresh weight (g) within thirty minutes of collection and used the Petiole smartphone application with calibration pad N°7 (Petiole LTD; https://petioleapp.com/) to measure leaf area (cm^2^). We then oven dried leaves (as outlined above) and measured dry weight (g). Leaf dry-matter content (LDMC; g/g) was measured as dry weight divided by fresh weight, and specific leaf area (SLA; cm^2^/g) was measured as leaf area divided by dry weight (Pérez-Harguindeguy et al., 2013). Leaves were then ground into a fine powder with a Retsch mixer mill MM 400 (Retsch, Haan, Germany) and analysed for percent carbon (%C), percent nitrogen (%N), and C:N with an Elementar vario EL cube CHNOS Elemental Analyzer (Elementar, Langenselbold, Germany), with a certified birch leaf reference standard (Elementar Microanalysis, Devon, UK). We did not obtain results from plants in the last experimental block (replicate 12) due to a technical issue during this analysis. As such, we analysed nutrient data from 214 of the 234 surviving plants (replicates 1–11).

### 2.4 | Statistical analyses

All analyses were conducted in R v4.0.3 (R Core Team, 2020) with RStudio v1.4.1103 (RStudio Team, 2021). We tested for differences between range-core and range-margin cohorts with a series of mixed effects models using the *lmer* function in the lmerTest R-package (Kuznetsova, Brockhoff, & Christensen, 2017), with source region as a fixed effect and maternal cohort nested within region as a random effect. Although maternal cohorts were clustered by collection site nested within region, we did not include this random effect as variation attributed to environmental differences among collection sites should be accounted for with our inclusion of propagule weight as a covariate to account for maternal investment (see detailed model descriptions below). For linear models, we assessed fixed effects with the *anova* function with default Type III SS, and we assessed random effects with the *ranova* function with likelihood ratio tests. For the generalised linear model, all effects were assessed with likelihood ratio tests. Refer to Table S1 in the Supporting Information for detailed summaries of each model described below.

First, we tested whether the weight of field-collected propagules, a measure of maternal investment, varied between source regions and among maternal cohorts nested within region with a linear mixed effects model. Next, for the establishment phase of the experiment (0–8 months), we tested for effects of source region and maternal cohort nested within region on seedling survival (binary response) with a binomial generalised linear mixed effects model, and on time to establishment, height at 8 months, and total growth at 8 months with linear mixed effects models. We included propagule weight as a covariate (fixed) in these models to account for variation in maternal investment and replicate planting tray as a (random) blocking factor to account for environmental variation within the greenhouse. We included time to establishment as an additional covariate (fixed) in the height and growth models because it proved influential for both response variables, independent of propagule weight (see Table S1).

For the subsequent growth phase of the experiment (8–24 months), we tested for effects of source region and maternal cohort nested within region on height, total growth, biomass, ratios of height/growth to biomass, plant stress (F_v_/F_m_), and leaf traits (leaf area, LDMC, SLA, %C, %N, and C:N) with linear mixed effects models. We log-transformed C:N for statistical analyses (Isles, 2020). We included plant size at the start of this phase of the experiment (i.e., total growth at 8 months) as a covariate (fixed) to account for variation in both propagule weight (measure of maternal investment) and time to establishment. In addition, we included replicate planting tray as a (random) blocking factor to account for environmental variation within the greenhouse. As height and total growth were measured at five time points, we first used repeated-measures models that included the effect of time (fixed) and the time × source region interaction (fixed) before analysing individual time points. We used the same approach for plant stress (F_v_/F_m_), which was measured at three time points.

Visual inspection of diagnostic plots for each model confirmed that linear models with a normal error distribution were suitable for all variables, except for survival that was assessed with a binomial error distribution. We did identify two large outliers for both SLA and log-transformed C:N, which were removed for analyses (see Table S1), although their inclusion did not qualitatively change the results described here. From each model, we calculated estimated marginal means for each source region in the emmeans R-package (Lenth, 2021). We also calculated marginal R^2^ (variability explained by fixed effects) and conditional R^2^ (variability explained by both fixed and random effects) for each model with the *r.squaredGLMM* function in the MuMIn R-package (Bartoń, 2020). Values for each model are presented in Table S1.

## 3 | RESULTS

### 3.1 | Maternal genotypes

All 20 maternal trees were genetically distinct, with a clear separation between range-core and range-margin genotypes (Figure 1C). Range-margin maternal trees exhibited greater genetic differences and greater clustering by collection site compared to those from the range core (Figure 1C).

### 3.2 | Greenhouse conditions

Greenhouse temperatures were relatively consistent with long-term averages at the Atlantic Florida range margin (Figure S1). Chilling temperatures (≤10°C) were experienced on 29 and 82 days during the first and second year of the experiment, respectively. The number of days ≤10°C during the second year was higher than what is generally experienced at the Atlantic Florida range margin (73 ± 1.5 days; Devaney et al., 2021). Sub-zero temperatures were experienced on only two days (2 – 3 Feb 2019), both during the second year of the experiment (min: −1 and −3°C, respectively). Greenhouse relative humidity (%) was 55.0 ± 6.3 (SD) across the experimental period, considerably lower than annual values at the Atlantic Florida range margin (76.6 ± 7.9; data from St Augustine Airport, obtained from https://www.ncdc.noaa.gov/data-access/land-based-station-data/land-based-datasets). Salinity (‰) within replicate trays was 14.0 ± 2.7 (SD) for the initial establishment phase of the experiment and was 18.6 ± 4.3 for the subsequent growth phase.

### 3.3 | Establishment phase (0 – 8 months)

Weight of field-collected propagules, a measure of maternal investment, varied among maternal cohorts (χ^2^(1) = 235.7, p < .001), with a mean increase of 98% from the cohort with the lightest to heaviest propagules (1.85 to 3.64 g) (Figure S2). Propagules from range-margin cohorts were heavier than those from range-core cohorts (F_1, 18_ = 7.7, p = .013), with a 23% increase in the estimated marginal mean (2.77 and 2.26 g, respectively), although there was considerable variation among range-margin cohorts (Figure S2).

A total of 529 of 600 planted propagules (88.2%) survived to establishment. Survival (χ^2^(1) = 31.2, p < .001), time to establishment (χ^2^(1) = 98.4, p < .001), height at 8 months (χ^2^(1) = 94.0, p < .001), and total growth at 8 months (χ^2^(1) = 122.7, p < .001) all varied among maternal cohorts (Figure 2). Survival ranged from only 40% within one range-core cohort to 100% within six range-margin cohorts (Figure 2A). Mean increases from the cohort with the lowest to highest values were 14% for establishment time (26.0 to 30.2 weeks) and 90% for both height and total growth (13.6 to 25.9 cm each) (Figure 2B, C). Range-margin cohorts survived in greater numbers (96%; 289 of 300 planted propagules) compared to range-core cohorts (80%; 240 of 300 planted propagules) (χ^2^(1) = 12.1, p = .005) (Figure 2A), and established faster (F_1, 18.9_ = 7.4, p = .014) with a 4% decrease in the estimated marginal mean (27.1 and 28.3 weeks) (Figure 2B). Several range-margin cohorts grew more than their range-core conspecifics over the first eight months, but there were two notable exceptions that were among the smallest plants in the experiment (cohort: GN4, GN5; Figure 2C). As a result, estimated marginal means were nearly identical between regions for both height (F_1, 20.5_ = 0.9, p = .356) and growth (F_1, 20.1_ = 0.3, p = .586) (Figure 2C). Propagule weight did not affect survival (χ^2^(1) = 1.2, p = .265) or establishment time (F_1, 466.3_ = 0.4, p = .525), but impacted height (F_1, 416.3_ = 52.5, p = .001) and growth (F_1, 407.2_ = 66.7, p = .001). Time to establishment also impacted height (F_1, 512.7_ = 305.5, p < .001) and growth (F_1, 494.8_ = 268.5, p < .001).

**FIGURE 2.**
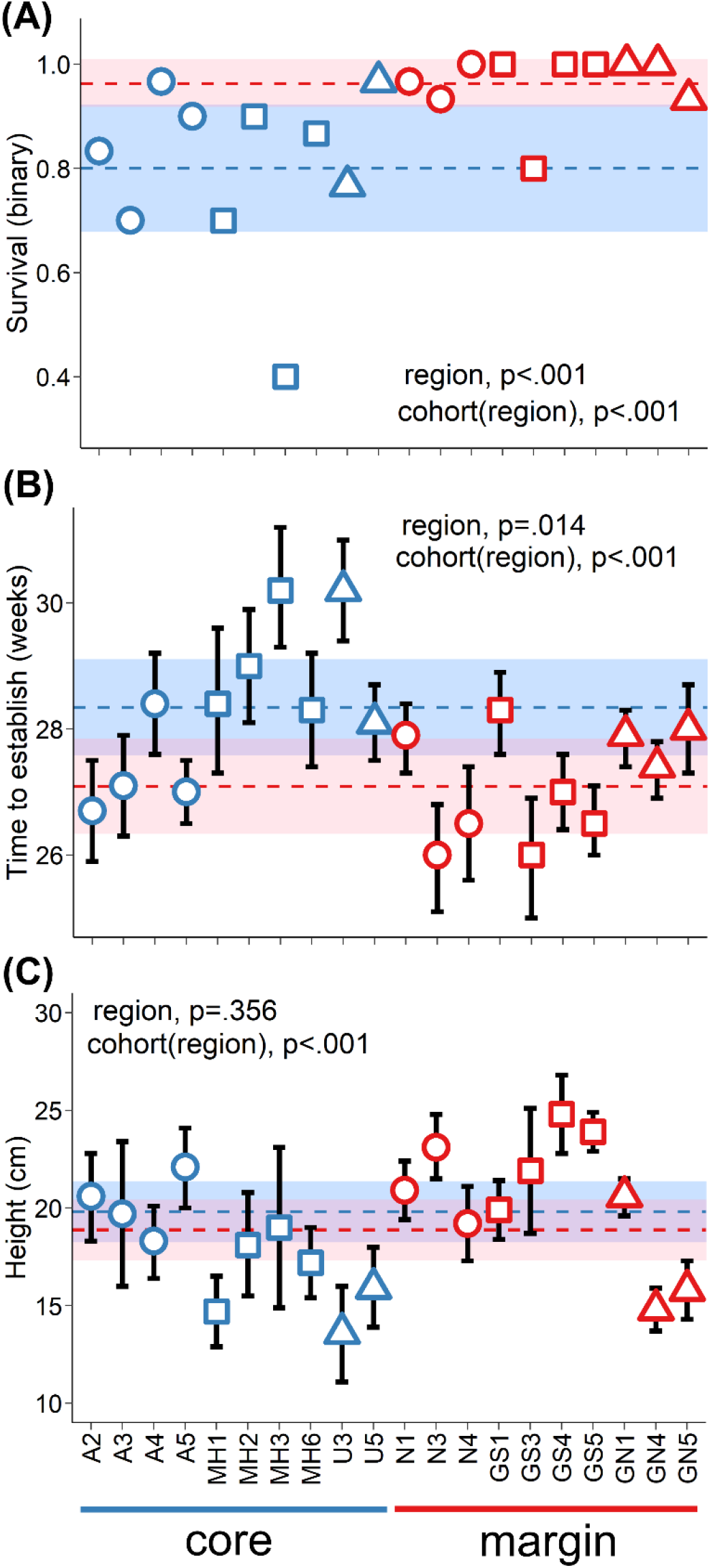
Range-margin cohorts (shown in red) (A) survived in greater numbers and (B) established faster, but (C) exhibited similar height, and total growth (*not shown*), at eight months compared to range-core cohorts (shown in blue). In the figure, different colour/shape combinations depict the six collection sites (refer to Figure 1 for geographical locations). Region-level estimated marginal means and 95% confidence intervals are shown with dashed lines and shaded areas in blue for the range core and in red for the range margin. Cohort-level means and 95% confidence intervals are calculated from the raw data.

### 3.4 | Subsequent growth phase (8 – 24 months)

A total of 234 of 240 transferred seedlings (97.5%) survived the subsequent 16 months of the experiment. The six mortalities consisted of three range-core and three range-margin plants.

We assessed plant stress with measurements of the maximum quantum yield of photosystem II (F_v_/F_m_) in December 2018, February 2019, and September 2019. Mean temperatures were 14.3°C (min / max: 9.8 / 18.7°C), 6.6°C (−0.7 / 16.8°C), and 20.4°C (13.3 / 34.8°C) during these measurement periods, respectively. We found that the effect of source region on plant stress varied temporally (time × source region: F_2, 666.1_ = 54.7, p < .001; Table S1), with the strongest effect of source region observed during the most stressful temperature conditions (February 2019). Maternal cohorts exhibited variation in F_v_/F_m_ at each time point (Dec 2018: χ^2^(1) = 6.8, p = .009; Feb 2019: χ^2^(1) = 18.5, p < .001; Sept 2019: χ^2^(1) = 13.3, p < .001), with mean increases from the cohort with lowest to highest F_v_/F_m_ of 15% (0.57 to 0.66), 78% (0.26 to 0.47), and 5% (0.73 to 0.76), respectively (Figure 3). Range-margin cohorts consistently had higher F_v_/F_m_ than range-core cohorts (Dec 2018: F_1, 18.6_ = 39.4, p < .001; Feb 2019: F_1, 18.7_ = 60.7, p < .001; Sept 2019: F_1, 18.9_ = 15.9, p = .001), with increases in estimated marginal means of 9% (0.64 and 0.59), 37% (0.41 and 0.30), and 1% (0.75 and 0.74) across the three time points, respectively (Figure 3). Total growth at 8 months impacted F_v_/F_m_ in December 2018 (F_1, 184.5_ = 8.4, p = .004) and February 2019 (F_1, 211.4_ = 12.9, p < .001), but not in September 2019 (F_1, 202.6_ = 1.4, p = .246).

**FIGURE 3.**
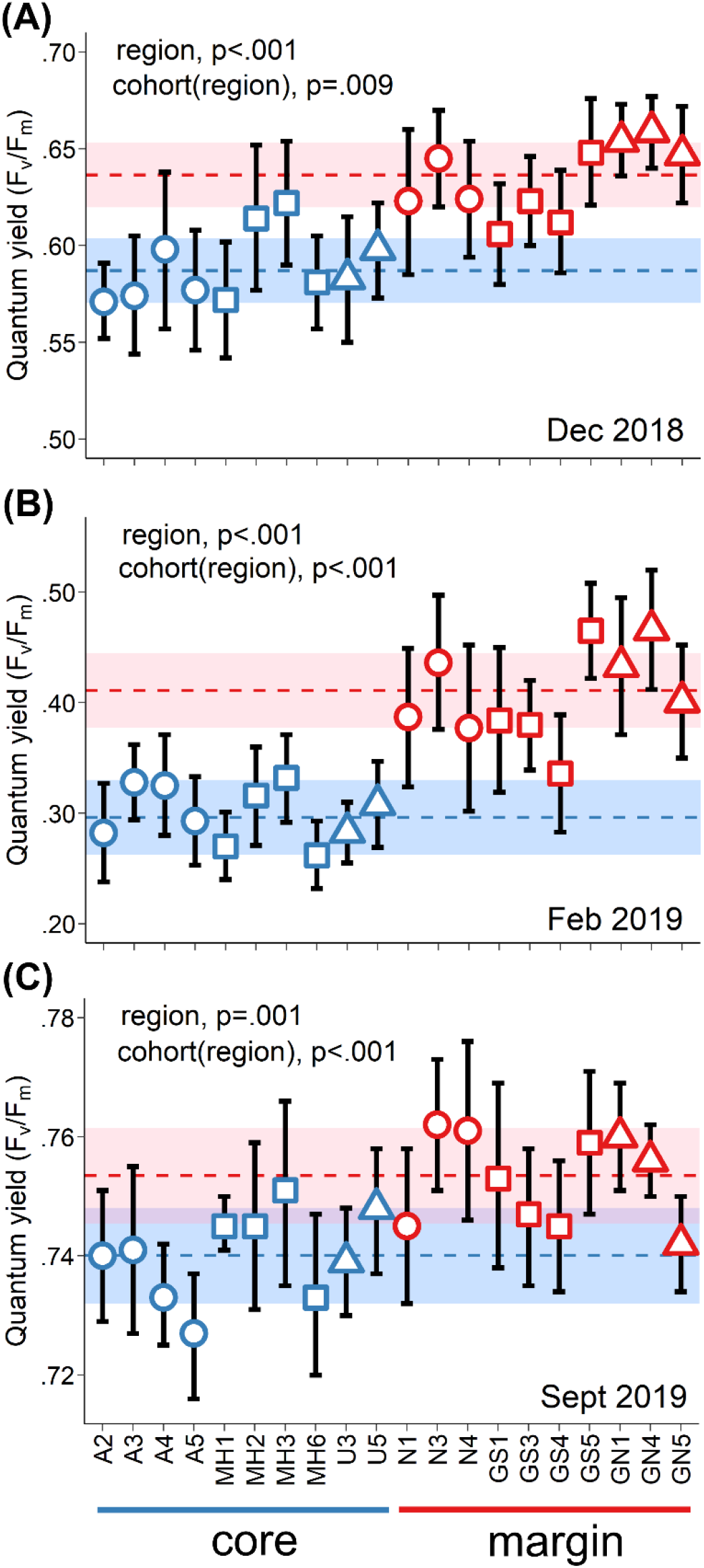
Range-margin cohorts (shown in red) were less stressed than range-core cohorts (shown in blue) in (A) December 2018, (B) February 2019 after consecutive nights of sub-zero temperatures, and (C) September 2019. Note that y-axes vary among panels. In the figure, different colour/shape combinations depict the six collection sites (refer to Figure 1 for geographical locations). Region-level estimated marginal means and 95% confidence intervals are shown with dashed lines and shaded areas in blue for the range core and in red for the range margin. Cohort-level means and 95% confidence intervals are calculated from the raw data.

We measured plant height and total growth at 10, 12, 14, 20, and 24 months post-planting. We found that the effect of source region on height varied temporally (time × source region: F_4, 1130.0_ = 6.6, p < .001); whereas, we found no temporal variation in the effect of source region on total growth (time × source region: F_4, 1130.0_ = 0.7, p = .625) (Table S1). Height varied among maternal cohorts at every time point (p < .001; Table S1). At 24 months, height (χ^2^(1) = 39.0, p < .001) and total growth (χ^2^(1) = 28.0, p < .001), varied among maternal cohorts, with mean increases from the cohort with lowest to highest values of 55% (36.7 to 56.9 cm) and 63% (47.2 to 77.1 cm), respectively (Figure 4A, B). As found at 8 months, the height of range-margin cohorts did not vary from those of range-core cohorts at 10 months (F_1, 18.7_ = 0.3, p = .582) or at 12 months (F_1, 18.5_ = 2.5, p = .134). However, starting at 14 months, range-margin cohorts were marginally shorter than range-core cohorts (F_1, 18.6_ = 4.5, p = .047) and this difference progressively became larger at 20 months (F_1, 18.4_ = 5.6, p = .029) and then at 24 months (F_1, 18.6_ = 7.5, p = .013), when we found a decrease in the estimated marginal mean of 8% (48.3 and 52.2 cm) (Figure 4A). In contrast, estimated marginal means for total growth at 24 months were nearly identical between regions (F_1, 18.6_ = 0.0, p = .844) (Figure 4B). As detailed for height/growth at 8 months, these patterns were partly shaped by two range-margin cohorts that were notable exceptions and among the smallest plants in the experiment (cohort: GN4, GN5; Figure 4A, B). Total growth at 8 months had a substantial impact on height at 24 months (F_1, 222.6_ = 237.5, p < .001) and total growth at 24 months (F_1, 226.5_ = 452.3, p < .001).

**FIGURE 4.**
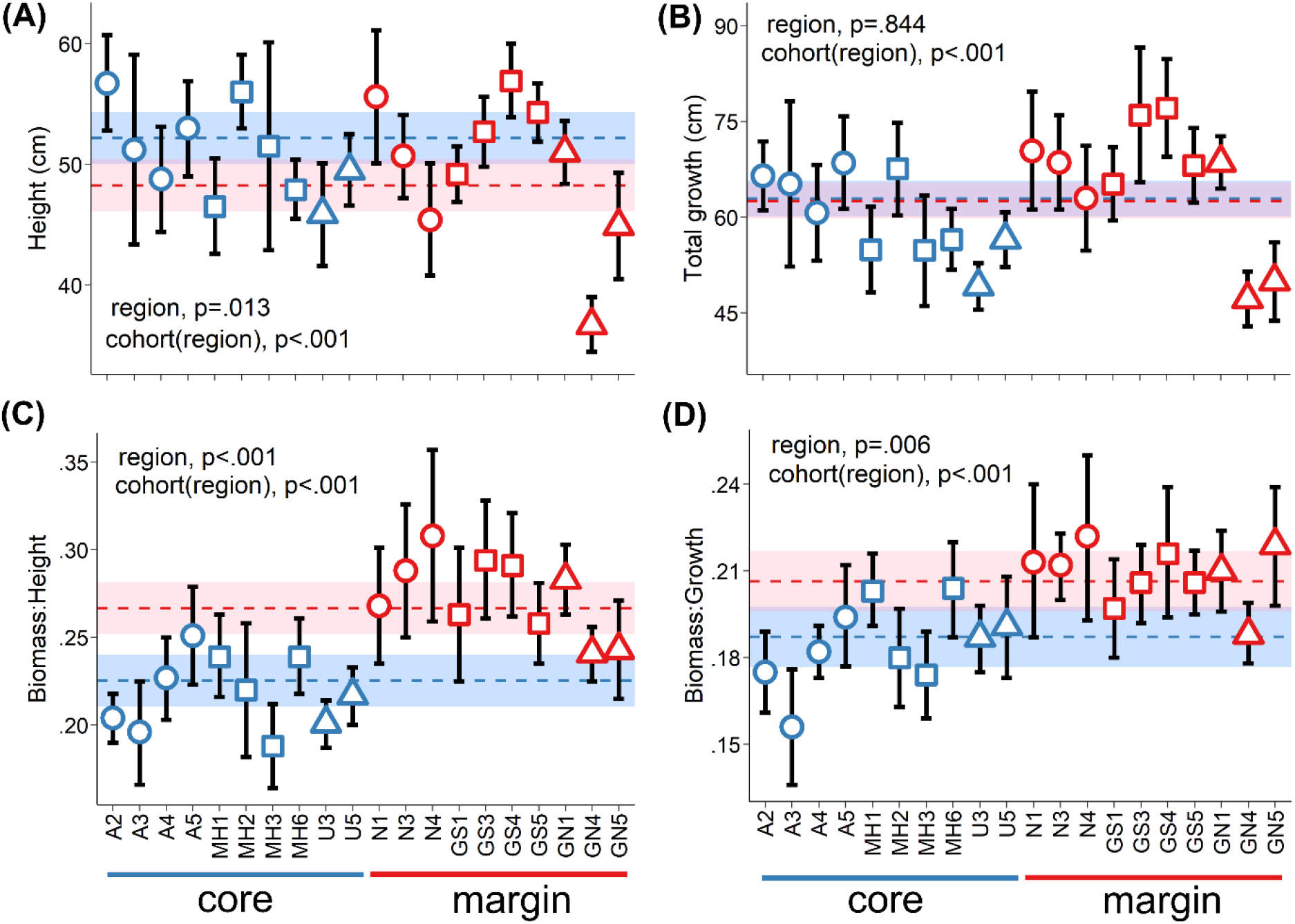
At two years development, range-margin cohorts (shown in red) were (A) shorter, but (B) exhibited similar total growth (height plus lateral growth), and accumulated a greater proportion of biomass (C) to height and (D) to growth compared to range-core cohorts (shown in blue). In the figure, different colour/shape combinations depict the six collection sites (refer to Figure 1 for geographical locations). Region-level estimated marginal means and 95% confidence intervals are shown with dashed lines and shaded areas in blue for the range core and in red for the range margin. Cohort-level means and 95% confidence intervals are calculated from the raw data.

Total biomass (χ^2^(1) = 35.9, p < .001) and the ratios of biomass to height (χ^2^(1) = 19.3, p < .001) and to growth (χ^2^(1) = 20.5, p < .001) all varied among maternal cohorts, with mean increases from the cohort with lowest to highest values of 87% (8.9 to 16.6 g), 63% (0.19 to 0.31), and 44% (0.16 to 0.23), respectively (Figure 4C, D; Figure S3). Range-margin cohorts accumulated more biomass (F_1, 18.5_ = 7.1, p = .015) and exhibited greater biomass to height (F_1, 18.3_ = 21.5, p < .001) and to growth (F_1, 18.1_ = 9.6, p = .006) compared to range-core cohorts, with increases in estimated marginal means of 11% (13.1 and 11.8 g), 17% (0.27 and 0.23), and 11% (0.21 and 0.19), respectively (Figure 4C, D; Figure S3). Range-margin cohorts tended to accumulate more biomass across each measured fraction (i.e., leaves, shoots, and roots), but region-level differences were only statistically-significant for leaves (F_1, 18.3_ = 10.8, p = .004) and roots (F_1, 18.7_ = 9.8, p = .006), not shoots (F_1, 18.6_ = 0.2, p = .704) (Figure S3). Again, these patterns were partly shaped by two smaller range-margin cohorts (cohort: GN4, GN5; Figure S3). Total growth at 8 months had a substantial impact on total biomass (F_1, 227.3_ = 422.8, p < .001) and impacted the ratios of biomass to height (F_1, 219.3_ = 97.0, p < .001) and to growth (F_1, 220.9_ = 25.9, p < .001).

Leaf area (χ^2^(1) = 39.8, p < .001), leaf dry-matter content (LDMC; χ^2^(1) = 25.8, p < .001), specific leaf area (SLA; χ^2^(1) = 14.6, p < .001), and log-transformed C:N (χ^2^(1) = 8.0, p = .005) all varied among maternal cohorts, with mean increases from the cohort with lowest to highest values of 62% (6.9 to 11.2 cm^2^), 15% (0.33 to 0.38 g/g), 26% (57.4 to 72.1 cm^2^/g), and 8% (3.39 to 3.67), respectively (Figure 5). Leaf area (F_1, 18.7_ = 0.0, p = .844) and LDMC (F_1, 18.3_ = 1.4, p = .251) did not vary between range-margin and range-core cohorts (Figure 5A, B). Instead, range-margin cohorts exhibited greater SLA (F_1, 19.0_ = 51.2, p < .001) and lower log-transformed C:N (F_1, 17.0_ = 12.9, p = .002) compared to range-core cohorts, with an increase in the estimated marginal mean of 12% (66.5 and 59.4 cm^2^/g) and a decrease of 3% (3.48 and 3.58), respectively (Figure 5C, D). Lower C:N in the leaves of range-margin cohorts was the product of greater %N (F_1, 16.9_ = 10.8, p = .004) and not changes in %C (F_1, 18.6_ = 0.8, p = .370) (Figure S4). Total growth at 8 months impacted LDMC (F_1, 220.3_ = 8.4, p = .004) and SLA (F_1, 210.0_ = 22.0, p < .001), but did not impact leaf area (F_1, 215.7_ = 0.0, p = .931) or log-transformed C:N (F_1, 179.1_ = 0.0, p = .923).

**FIGURE 5.**
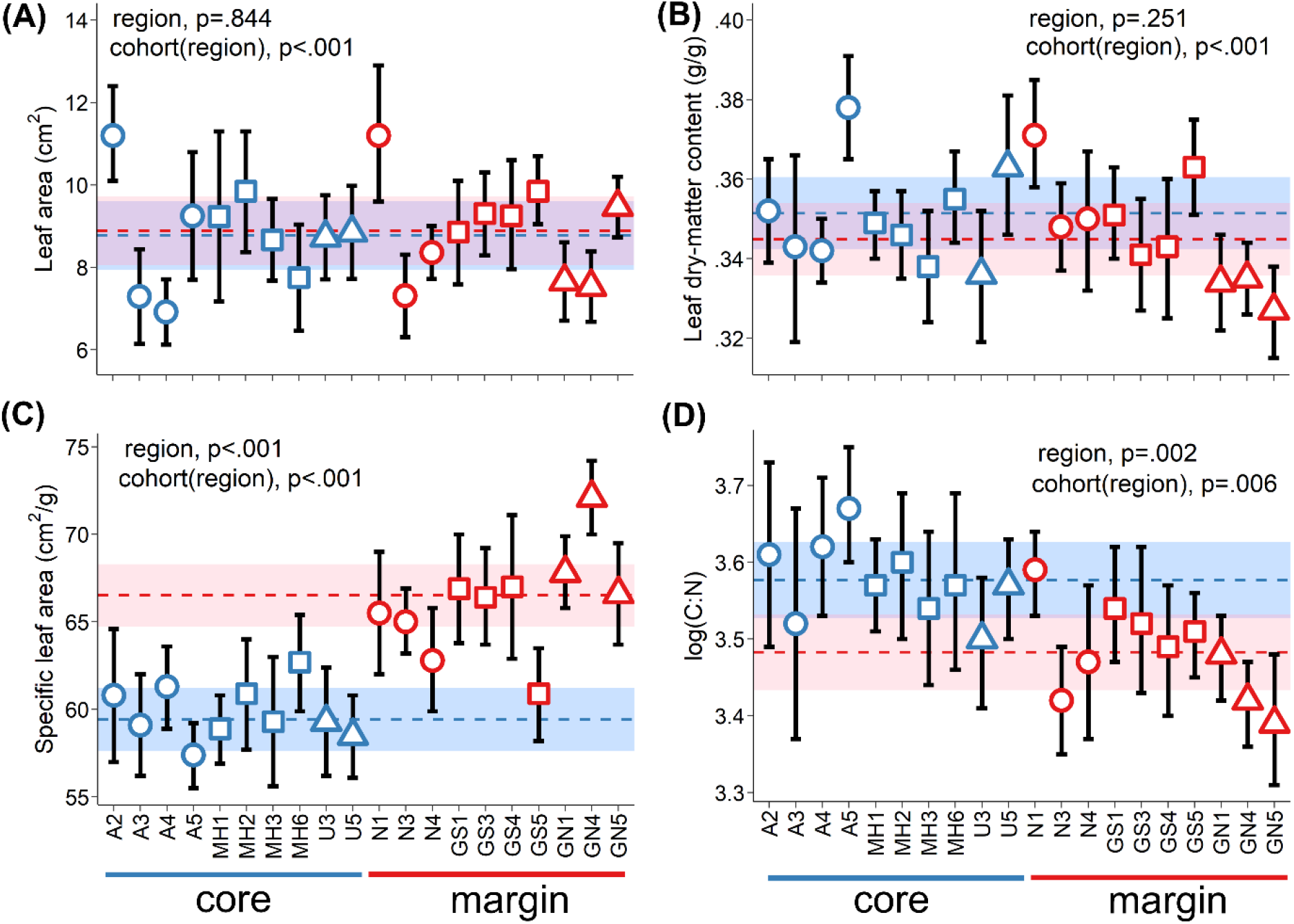
Range-margin cohorts (shown in red) produced leaves of similar (A) size and (B) leaf dry-matter content (LDMC), but with (C) greater specific leaf area (SLA) and (D) lower C:N compared to range-core cohorts (shown in blue). In the figure, different colour/shape combinations depict the six collection sites (refer to Figure 1 for geographical locations). Region-level estimated marginal means and 95% confidence intervals are shown with dashed lines and shaded areas in blue for the range core and in red for the range margin. Cohort-level means and 95% confidence intervals are calculated from the raw data.

## 4 | DISCUSSION

In this study, we confirmed that there is a genetic basis to adaptive trait shifts towards an expanding range margin of a mangrove foundation species (*Avicennia germinans*). Maternal cohorts from the northern Atlantic Florida range margin consistently outperformed those from the southern range core under annual temperatures analogous to range-margin conditions in a two-year greenhouse common garden experiment. Our findings suggest that genetically-based phenotypic differences better enable these range-margin mangroves to thrive under their stressful conditions and may facilitate further range expansion with climate change. In addition, our documentation of substantial adaptive trait variation among maternal cohorts of an ecologically-important mangrove species should help inform future mangrove restoration initiatives.

### 4.1 | Range-margin mangroves outperform range-core conspecifics

Species at their range margins are often genetically distinct from range-core conspecifics and may also exhibit adaptive shifts in morphology, reproductive strategies, and stress tolerance to facilitate establishment and survival in their marginal environment (Chuang & Peterson, 2016). The twenty *A. germinans* maternal trees sampled for this experiment exhibited a clear distinction between range-margin and range-core genotypes, consistent with population-level genetic differences along Atlantic Florida (Kennedy, Preziosi, et al., 2020). Also, we found that cohorts of field-collected propagules from range-margin trees planted in a common garden exhibited clear advantages over range-core cohorts during their critical establishment phase and under stressful winter conditions. In support of our first prediction, range-margin cohorts not only survived in greater numbers, but these survivors established, on average, more than a week earlier than range-core cohorts. We then observed that all plants exhibited signs of stress [i.e., suboptimal values of quantum yield (F_v_/F_m_)] under winter chilling and sub-zero temperatures, a ubiquitous plant response to winter conditions (Oliveira & Peñuelas, 2005). Yet, in support of our second prediction, range-margin cohorts exhibited higher F_v_/F_m_ under chilling stress and the difference between range-margin and range-core cohorts was even greater when temperatures dropped below 0°C. These differences suggest that range-margin cohorts were better able to maintain photosynthetic efficiency under winter conditions, and that this ability was more pronounced under more extreme conditions often experienced at the Atlantic Florida range margin, but not within the range core. Close to optimal F_v_/F_m_ (cohort means ranged from 0.73 to 0.76), with minimal differences between source regions, under more benign summer temperatures suggests that all plants subsequently recovered photosynthetic efficiency. Similar values approaching 0.8 are documented in *A. germinans* seedlings under optimal light and salinity conditions (Dangremond, Feller, & Sousa, 2015).

We found mixed support for our third prediction as greater stress tolerance in range-margin cohorts did translate into greater biomass accumulation, but not into greater growth. Instead, range-core cohorts gradually grew taller as the experiment progressed, while total growth (height plus length of lateral shoots) remained nearly identical between source regions throughout the experiment. In other words, over time, range-margin cohorts invested more into lateral versus vertical growth. Adult *A. germinans* at the Atlantic Florida range margin demonstrate this same pattern as they grow wider rather than taller (Chapman et al., 2021). Range-margin cohorts also accumulated a greater proportion of biomass relative to their size. This shift towards a greater investment into biomass over height may reflect adaptation to novel conditions within the harsh range-margin environment, analogous to responses across elevation gradients (Parker, Rodriguez, & Loik, 2003). At poleward range margins, shorter *A. germinans* would be less impacted by cold events due to warmer temperatures closer to the soil surface (Osland et al., 2019) and to protection offered by salt marsh vegetation (Pickens, Sloey, & Hester, 2019). Increased height would also not be as beneficial for developing range-margin mangroves in terms of greater access to sunlight as even juvenile trees can outcompete the surrounding low-stature salt marsh for light (Guo, Zhang, Lan, & Pennings, 2013). In contrast, increased height would be essential for range-core mangroves attempting to reach sunlight within closed canopy forests (Krauss et al., 2008). It is important to note that the patterns outlined here were partly shaped by two range-margin cohorts that presented obvious exceptions in terms of growth. These cohorts, both from the most northern collection site, were consistently among the smallest plants in the experiment. However, despite their small stature, these plants were not underperforming as they exhibited clear advantages over range-core cohorts at many other measured traits, including greater stress tolerance, greater proportion of biomass to height/growth, and greater resource acquisition (see discussion in the next paragraph). A reciprocal transplant experiment with planting sites in both the range core and margin could assess whether reduced height represents local adaptation within range-margin *A. germinans*.

Our fourth prediction was not supported as range-margin cohorts did not exhibit more conservative leaf traits (i.e., smaller, increased dry-matter content, reduced specific leaf area), which suggests that previous field documentation of systematic shifts in these particular traits among populations of Atlantic Florida *A. germinans* (Cook-Patton et al., 2015; Kennedy, Preziosi, et al., 2020) may be the product of trait plasticity in response to environmental variation. Instead, in the common garden, we found that range-margin cohorts produced leaves of similar size and leaf dry-matter content to those of range-core cohorts, but with increased specific leaf area and lower C:N due to greater nitrogen content. These differences within range-margin cohorts may reflect leaf development under a less stressed state (Poorter, Niinemets, Poorter, Wright, & Villar, 2009) and metabolic adjustments common in plants adapted to cold temperatures (Janská, Maršík, Zelenková, & Ovesná, 2010; Woods et al., 2003). These differences are also consistent with a greater ability among range-margin cohorts to capture light and nutrient resources, further supported by their greater accumulation of leaf and root biomass compared to range-core cohorts. Plastic shifts towards increased specific leaf area and root growth are found in *A. germinans* seedlings under limited resource availability to maximise resource acquisition (McKee, 1995). Here, we found that similar genetically-based shifts may also occur along a relatively narrow transition from mangrove range core to margin (27.5 – 30.0°N), presumably to compensate for a shorter growing season and reduced light quality at higher latitude (Spence & Tingley, 2020). An analogous genetically-based shift towards greater resource acquisition is also documented across greater geographic distance (0 – 28°S) between range core and margin populations of *A. schaueriana*, a closely-related congener (Cruz et al., 2019).

### 4.2 | Genetic basis to trait variation in range-margin mangroves

Range margins may foster unique genetic adaptations that enable species to persist under extreme climatic conditions and that can dictate future responses to climate change (Rehm et al., 2015). Here, we demonstrated a genetic basis to adaptative trait shifts within a mangrove towards its expanding Atlantic Florida range margin, with evidence of greater survival during initial establishment, greater stress tolerance over winter, greater biomass accumulation, and greater resource acquisition among range-margin cohorts. Although still limited, growing evidence supports genetically-based adaptive shifts in chilling tolerance (Markley, McMillan, & Thompson Jr, 1982; Short, Chen, & Wee, 2021), freezing tolerance (Hayes et al., 2020), resource acquisition (Cruz et al., 2019), and precocious reproduction (Dangremond & Feller, 2016) towards cold-sensitive mangrove distributional margins in Brazil, China, and the USA. In addition, evidence for selection along climatic gradients is found across mangrove distributions in Brazil (Cruz et al., 2020; Da Silva et al., 2021). Therefore, despite the immense trait plasticity within mangroves that enables their proliferation across highly variable environments (Feller et al., 2010), trait evolution may also be a common phenomenon in these systems, in particular towards range margins where selection pressures are inherently at their highest. Multiple interacting processes could drive this change, including selective mass mortality, genetic drift, and spatial sorting (Nadeau & Urban, 2019), as well as epigenetic changes (Robertson & Richards, 2015). A broader understanding of the processes driving these adaptive shifts could be achieved with further research that evaluates trait changes towards multiple mangrove range margins defined by distinct climatic thresholds and colonisation histories (e.g., Bardou, Parker, Feller, & Cavanaugh, 2021).

Our findings also provide insight into how an ecologically-important mangrove species (*A. germinans*) may respond to climate change at its poleward range margin. Phenotypic differences outlined above present clear advantages for range-margin over range-core genotypes in terms of proliferation within currently occupied range-margin sites and colonisation of more poleward areas. Mangrove expansion is forecast along Atlantic Florida as freeze events become less common (Cavanaugh et al., 2019, 2015), but poleward expansion of *A. germinans* along this coastline may be restricted to periods following extreme storm events that provide new recruits almost exclusively from range-margin sources (Kennedy, Dangremond, et al., 2020). Hence, an expanding gene pool with a greater representation of range-margin genotypes, that are better able to thrive under the climatic extremes beyond the current mangrove distribution, will presumably facilitate future *A. germinans* range expansion. This transition from salt marsh to mangrove dominance will inevitably have wide-reaching effects on these coastal ecosystems, including increased carbon storage, greater sediment accretion in response to sea level rise, enhanced storm protection, and reduced habitat availability for certain fauna that require open vegetation (Doughty et al., 2016; Kelleway et al., 2017; Osland et al., 2018; Simpson, Stein, Osborne, & Feller, 2019).

### 4.3 | Considerations and Next steps

Offspring may exhibit phenotypic differences as a result of several factors, specifically the genetic makeup of their parents, their growing environment, and maternal effects that are shaped by both maternal genetics and maternal environment (Wolf & Wade, 2009). In this research, we monitored the development of field-collected propagules in a single greenhouse environment. Therefore, differences observed among maternal cohorts are the product of parental genetics and maternal effects. We genotyped maternal trees, but lack information on the unique genotypes of each individual plant within this experiment. As a result, some variation within maternal cohorts will be attributed to differences in pollination (i.e., proportions of progeny that are selfed, outcrossed full-siblings, and outcrossed half-siblings). Geographical variation in mating system, however, should not have systematically impacted our region-level results as there is not a systematic change in outcrossing rates along our range core to margin sampling gradient (Kennedy et al., 2021).

We found that field-collected propagules from range-margin maternal trees were, on average, heavier than those from range-core trees, consistent with previous documentation of greater propagule weight towards the Atlantic Florida range margin for *A. germinans* (Nathan, 2020) and of greater propagule size for the co-occurring mangrove, *Rhizophora mangle* (Dangremond & Feller, 2016). Propagule size is often influenced by maternal environment and is a common proxy for maternal effects. Greater maternal investment into offspring can facilitate species expansion (Estrada, Wilson, NeSmith, & Flory, 2016), although the strength of environmentally-induced maternal effects in plants generally declines as offspring age (Maruyama et al., 2016). Propagule weight, and subsequently total growth at 8 months, both proved highly influential in terms of growth and biomass accumulation across our two-year experiment. However, our observations of phenotypic differences among maternal cohorts were not merely shaped by variation in maternal investment. After accounting for this variation, we still observed significant effects of source region on height and biomass. In addition, greater propagule weight had no discernible impact on survival or establishment time and subsequent total growth at 8 months had a limited impact (compared to source region) on stress tolerance and most leaf traits.

Controlled common garden experiments can determine whether there is a genetic basis to phenotypic differences within a species, but inherently lack the reality and complexity of natural field conditions. Our greenhouse experiment demonstrates that range-margin *A. germinans* maternal cohorts may be better suited to thrive under stressful temperature conditions analogous to those at the Atlantic Florida range margin over the first two years of their development. However, in addition to temperature, multiple interacting abiotic and biotic factors will influence the establishment, survival, and growth of these range-margin mangroves (Rogers & Krauss, 2019). Longer-term *in situ* common gardens are, therefore, a logical next step to better predict how these coastal foundation species will respond to climate change. Although challenging because of long generation times, networks of common gardens have provided a wealth of knowledge regarding how forest trees have adapted to different environments and how they may respond to changing environmental conditions (Alberto et al., 2013). A series of common gardens both at and beyond mangrove range margins could further our understanding of the long-term fitness and persistence of these mangroves and of the factors that may limit or facilitate further range expansion.

### 4.4 | Implications for mangrove rehabilitation and restoration

Initiatives to rehabilitate and restore degraded coastal ecosystems are growing in number (Waltham et al., 2020). Mangrove foundation species are a central component of many such initiatives because of the ecosystem services they provide (Friess et al., 2019). Restoration-focused experimental research demonstrates that intraspecific genetic and phenotypic variation within coastal foundation species can influence their survival and productivity, as well as ecosystem service provision (Bernik, Pardue, & Blum, 2018; Hughes, 2014; Plaisted, Novak, Weigel, Klein, & Short, 2020). Yet, only one study has documented similar quantitative data on adaptive trait variation within mangroves at the level that replanting occurs (i.e., propagules collected from maternal trees). Proffitt & Travis (2010) found that survival, growth, and age to reproduction varied among maternal cohorts of red mangrove (*Rhizophora mangle*) and that these patterns differed between two intertidal settings. Although mangrove replanting may often not be necessary (Lewis, 2005) and not a viable alternative at high-stress range margins (Macy, Osland, Cherry, & Cebrian, 2021), results from Proffitt & Travis (2010) and our documentation of substantial differences among mangrove maternal cohorts in survival, stress tolerance, growth, and biomass accumulation (key success criteria for rehabilitation and restoration projects) highlight how source selection could influence the outcome of initiatives where mangrove replanting is needed.

Clear advantages exhibited by range-margin cohorts grown under temperatures analogous to range-margin conditions could be viewed as support for using local sources, or sources with similar environmental conditions, in restoration projects (Bucharova et al., 2017). Records of propagule source and basic monitoring data on phenotypic variation within growth nurseries could help inform source selection and potentially improve replanting success. However, much more work is needed to understand how the genetic background of propagules used for replanting may influence the responses of these developing plants to the multiple interacting stressors common in mangrove systems (e.g., salinity, inundation, herbivory, irradiation) (Krauss et al., 2008). In addition, genetic variation within restoration plantings could shape the associated communities of organisms that colonise and inhabit these areas (Breed et al., 2018), with evidence that mangrove maternal genotype can influence soil microbial communities (Craig, Kennedy, Devlin, Bardgett, & Rowntree, 2020) and that genetic differences among mangrove hosts can correlate with the composition of endophytic fungal communities (Kennedy, Antwis, Preziosi, & Rowntree, *accepted*). Embedding *in situ* common garden experiments (as described in the previous section) into larger adaptive management experiments (Ellison, Felson, & Friess, 2020) could begin to uncover how intraspecific genetic variation may impact mangrove restoration and within which contexts these effects are most influential.

## ACKNOWLEDGEMENTS

This research was funded by a Manchester Metropolitan University studentship to J.P.K. Many thanks to IC Feller and R Feller (and their cast iron skillet) and the Smithsonian Marine Station at Fort Pierce, Florida for logistical support during fieldwork, to C Jameson, CM Kennedy, W Potdevin, and J Sammy for assistance with experiment set-up and growth measurements, to R Solymosi for R-programming help with monitoring data, to M da Silva and G Mori for sharing an R-script for climatic data analysis, to the Manchester Metropolitan University technical staff, in particular C Dean and D McKendry, for assistance throughout this project, and to the University of Manchester Genomic Technologies Core Facility and F Combe for fragment analysis. As always, thank you to A Jara Cavieres, C Kennedy, and M Kennedy for unconditional support and big smiles.

## AUTHOR CONTRIBUTIONS

J.P.K., G.N.J., R.F.P, and J.K.R. conceived the ideas and designed methodology. J.P.K. collected and analysed the data, and led the writing of the manuscript. All authors contributed critically to the drafts and gave final approval for publication.

## DATA AVAILABILITY STATEMENT

All data presented in this article will be publicly available on figshare.

## SUPPORTING INFORMATION

Additional supporting information may be found online in the Supporting Information section.

