## Supplemental Information for "Genetically-based adaptive trait shifts at an expanding mangrove range margin"

### TABLE OF CONTENTS


| Appendix | Page |
|---|---|
| Figure S1 | 2 |
| Figure S2 | 3 |
| Figure S3 | 4 |
| Figure S4 | 5 |
| Table S1 | 6–10 |

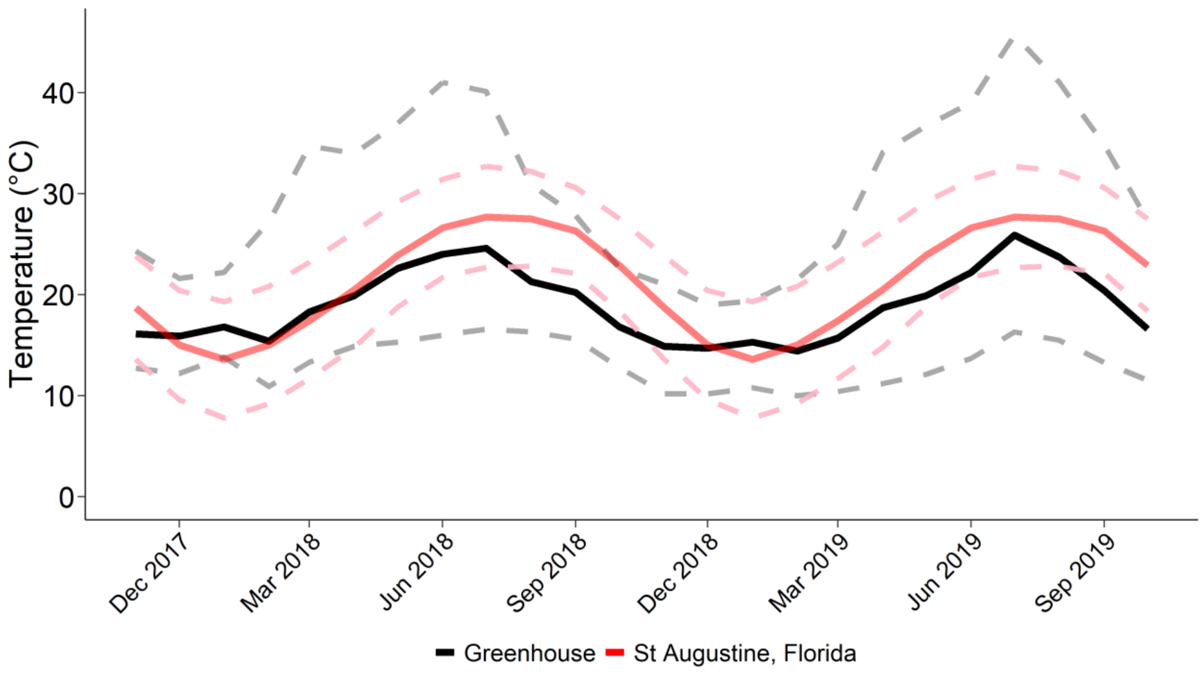


**FIGURE S1**. Greenhouse temperatures were relatively consistent with long-term (1981-2010) averages at the Atlantic Florida range margin (St Augustine, Florida; data from: https://www.ncdc.noaa.gov/cdo-web/datatools). Solid lines show monthly mean temperatures and dashed lines show monthly maximum and minimum temperatures.

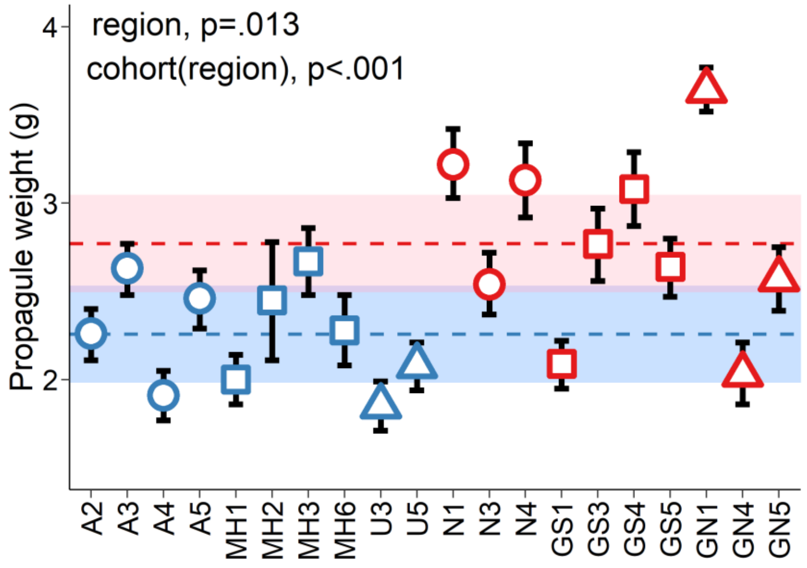


**FIGURE S2**. Field-collected propagules from range-margin cohorts (shown in red) were heavier than those from range-core cohorts (shown in blue), although considerable variation was found among range-margin cohorts. In the figure, different colour/shape combinations depict the six collection sites (refer to Figure 1 in the main text for geographical locations). Region-level estimated marginal means and 95% confidence intervals are shown with dashed lines and shaded areas in blue for the range core and in red for the range margin. Cohort-level means and 95% confidence intervals are calculated from the raw data.

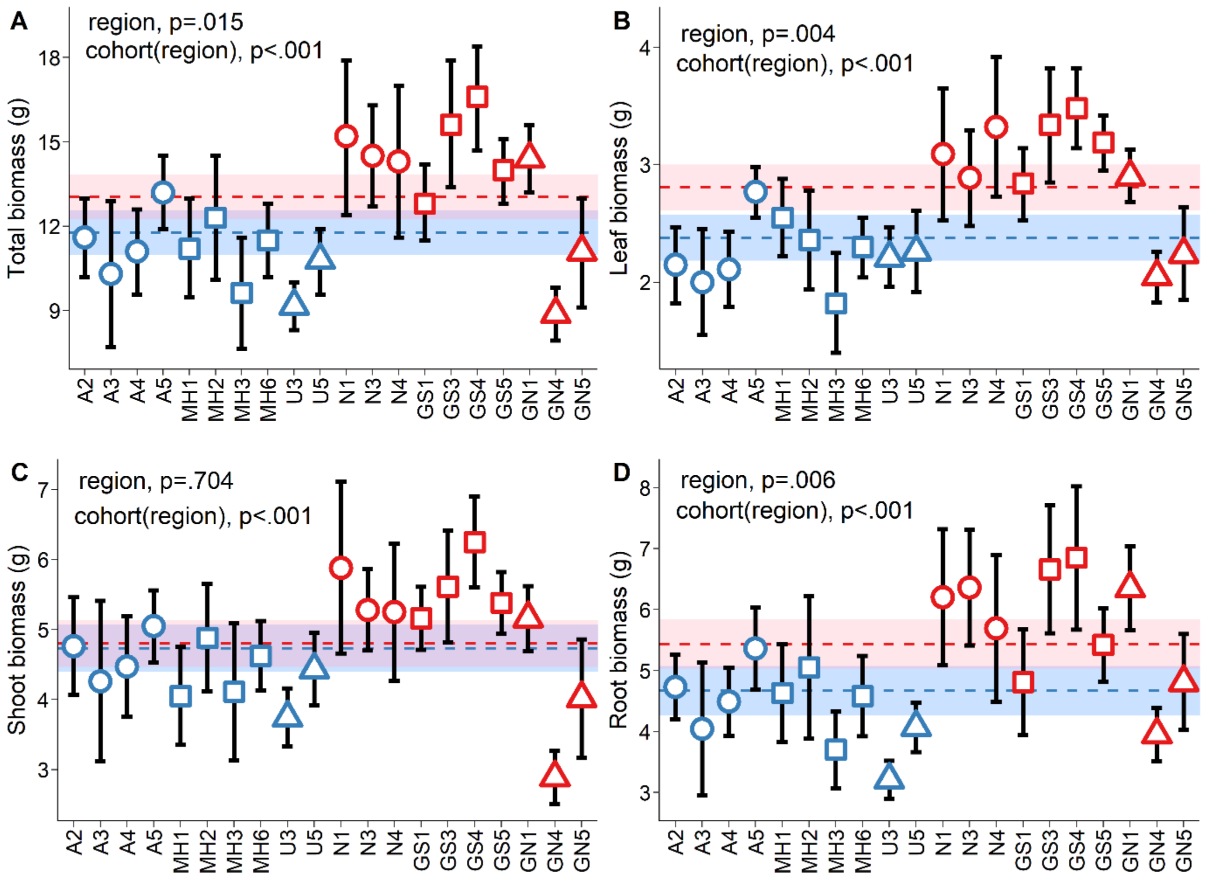


**FIGURE S3**. (A) Range-margin cohorts (shown in red) accumulated more biomass than range-core cohorts (shown in blue). Range-margin cohorts tended to accumulate more biomass across each measured fraction, (B) leaves, (C) shoots, and (D) roots, but region-level differences were only statistically-significant for leaves and roots. In the figure, different colour/shape combinations depict the six collection sites (refer to Figure 1 in the main text for geographical locations). Region-level estimated marginal means and 95% confidence intervals are shown with dashed lines and shaded areas in blue for the range core and in red for the range margin. Cohort-level means and 95% confidence intervals are calculated from the raw data.

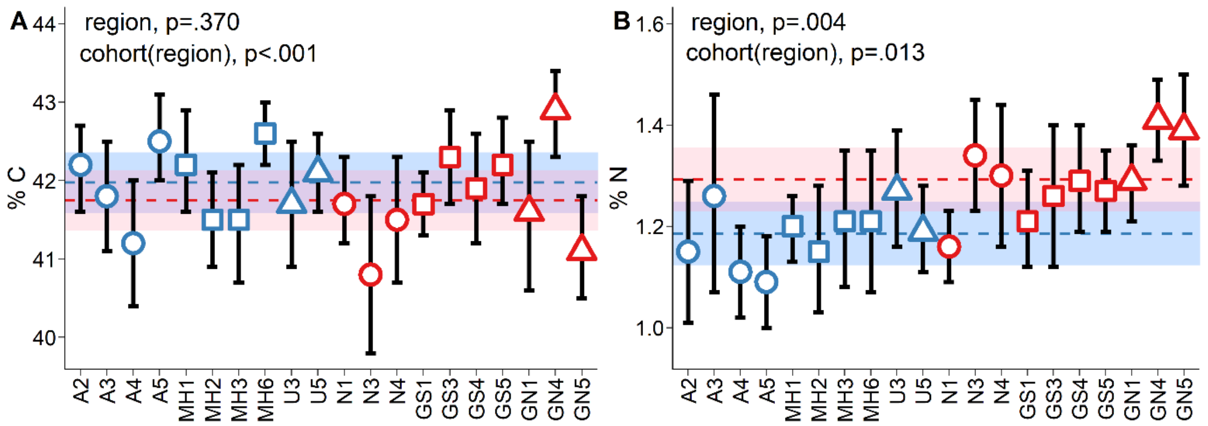


**FIGURE S4**. Range-margin cohorts (shown in red) produced leaves with (A) similar percent carbon, but (B) greater percent nitrogen compared to range-core cohorts (shown in blue). In the figure, different colour/shape combinations depict the six collection sites (refer to Figure 1 in the main text for geographical locations). Region-level estimated marginal means and 95% confidence intervals are shown with dashed lines and shaded areas in blue for the range core and in red for the range margin. Cohort-level means and 95% confidence intervals are calculated from the raw data.

**TABLE S1.** Response variables, predictor variables, and statistical results for the mixed effects models presented in the manuscript. $R^2_m$, variability explained by fixed effects; $R^2_c$, variability explained by fixed and random effects; Model, model structure used for analysis. Bold values indicate statistical significance ($p < 0.05$). [a] Two large outliers were removed to meet model assumptions. Results were equivalent when these data points were included.

| **Response** | **Predictors** | **Results** | **p-value** | **$R^2_m$** | **$R^2_c$** | **Model** |
|---|---|---|---|---|---|---|
| Propagule weight | Region | $F_{1, 18} = 7.7$ | **0.013** | 0.14 | 0.49 | lmer(prop_weight ~ region + (1 \| cohort)) |
| | Cohort(Region) | $\chi^2$ (1) = 235.7 | **< 0.001** | | | |
| Survival | Propagule weight | $\chi^2$ (1) = 1.2 | 0.265 | 0.21 | 0.53 | glmer(germ ~ prop_weight + region + (1 \| cohort) + (1 \| rep1), family = binomial()) |
| | Region | $\chi^2$ (1) = 12.1 | **<0.001** | | | |
| | Cohort(Region) | $\chi^2$ (1) = 31.2 | **< 0.001** | | | |
| | Block1 | $\chi^2$ (1) = 11.0 | **0.001** | | | |
| Time to establishment | Propagule weight | $F_{1, 466.3} = 0.4$ | 0.525 | 0.08 | 0.50 | lmer(time_germ ~ prop_weight + region + (1 \| cohort) + (1 \| rep1)) |
| | Region | $F_{1, 18.9} = 7.4$ | **0.014** | | | |
| | Cohort(Region) | $\chi^2$ (1) = 98.4 | **< 0.001** | | | |
| | Block1 | $\chi^2$ (1) = 116.3 | **< 0.001** | | | |
| Height at 8 months | Propagule weight | $F_{1, 416.3} = 52.5$ | **0.001** | 0.47 | 0.66 | lmer(height1 ~ prop_weight + time_germ + region + (1\|cohort) + (1 \| rep1)) |
| | Time to establishment | $F_{1, 512.7} = 305.5$ | **<0.001** | | | |
| | Region | $F_{1, 20.5} = 0.9$ | 0.356 | | | |
| | Cohort(Region) | $\chi^2$ (1) = 94.0 | **<0.001** | | | |
| | Block1 | $\chi^2$ (1) = 40.2 | **<0.001** | | | |
| Total growth at 8 months | Propagule weight | $F_{1, 407.2} = 66.7$ | **0.001** | 0.47 | 0.66 | lmer(grow1 ~ prop_weight + time_germ + region + (1 \| cohort) + (1 \| rep1)) |
| | Time to establishment | $F_{1, 494.8} = 268.5$ | **<0.001** | | | |
| | Region | $F_{1, 20.1} = 0.3$ | 0.586 | | | |
| | Cohort(Region) | $\chi^2$ (1) = 122.7 | **<0.001** | | | |
| | Block1 | $\chi^2$ (1) = 28.5 | **<0.001** | | | |

| | | | | | | |
|---|---|---|---|---|---|---|
| Quantum yield (repeated measures) | Growth at 8 months | $F_{1, 488.4} = 15.8$ | **<0.001** | 0.91 | 0.92 | lmer(CF ~ grow1 + time*region + (1 \| cohort) + (1 \| rep2)) |
| | Time | $F_{2, 666.1} = 3861.0$ | **<0.001** | | | |
| | Region | $F_{1, 18.6} = 60.8$ | **<0.001** | | | |
| | Time*Region | $F_{2, 666.1} = 54.7$ | **<0.001** | | | |
| | Cohort(Region) | $\chi^2$ (1) = 22.3 | **<0.001** | | | |
| | Block2 | $\chi^2$ (1) = 39.8 | **<0.001** | | | |
| Quantum yield (Dec. 2018) | Growth at 8 months | $F_{1, 184.5} = 8.4$ | **0.004** | 0.22 | 0.43 | lmer(DEC ~ grow1 + region + (1 \| cohort) + (1 \| rep2)) |
| | Region | $F_{1, 18.6} = 39.4$ | **<0.001** | | | |
| | Cohort(Region) | $\chi^2$ (1) = 6.8 | **0.009** | | | |
| | Block2 | $\chi^2$ (1) = 31.0 | **<0.001** | | | |
| Quantum yield (Feb. 2019) | Growth at 8 months | $F_{1, 211.4} = 12.9$ | **<0.001** | 0.34 | 0.62 | lmer(FEB ~ grow1 + region + (1 \| cohort) + (1 \| rep2)) |
| | Region | $F_{1, 18.7} = 60.7$ | **<0.001** | | | |
| | Cohort(Region) | $\chi^2$ (1) = 18.5 | **<0.001** | | | |
| | Block2 | $\chi^2$ (1) = 64.8 | **<0.001** | | | |
| Quantum yield (Sept. 2019) | Growth at 8 months | $F_{1, 202.6} = 1.4$ | 0.246 | 0.11 | 0.47 | lmer(CF8 ~ grow1 + region + (1 \| cohort) + (1 \| rep2)) |
| | Region | $F_{1, 18.9} = 15.9$ | **0.001** | | | |
| | Cohort(Region) | $\chi^2$ (1) = 13.3 | **<0.001** | | | |
| | Block2 | $\chi^2$ (1) = 59.9 | **<0.001** | | | |
| Height (repeated measures) | Growth at 8 months | $F_{1, 1128.8} = 1505.4$ | **<0.001** | 0.76 | 0.82 | lmer(height ~ grow1 + time*region + (1 \| cohort) + (1 \| rep2)) |
| | Time | $F_{4, 1130.0} = 624.2$ | **<0.001** | | | |
| | Region | $F_{1, 18.2} = 5.0$ | **0.039** | | | |
| | Time*Region | $F_{4, 1130.0} = 6.6$ | **<0.001** | | | |
| | Cohort(Region) | $\chi^2$ (1) = 231.1 | **<0.001** | | | |
| | Block2 | $\chi^2$ (1) = 41.1 | **<0.001** | | | |

| | | | | | | |
|---|---|---|---|---|---|---|
| Height at 10 months | Growth at 8 months | $F_{1, 215.6} = 578.9$ | **<0.001** | 0.74 | 0.80 | lmer(height2 ~ grow1 + region + (1 \| cohort) + (1 \| rep2)) |
| | Region | $F_{1, 18.7} = 0.3$ | 0.582 | | | |
| | Cohort(Region) | $\chi^2$ (1) = 16.2 | **<0.001** | | | |
| | Block2 | $\chi^2$ (1) = 5.6 | **0.018** | | | |
| Height at 12 months | Growth at 8 months | $F_{1, 210.6} = 341.1$ | **<0.001** | 0.62 | 0.69 | lmer(height3 ~ grow1 + region + (1 \| cohort) + (1 \| rep2)) |
| | Region | $F_{1, 18.5} = 2.5$ | 0.134 | | | |
| | Cohort(Region) | $\chi^2$ (1) = 14.0 | **<0.001** | | | |
| | Block2 | $\chi^2$ (1) = 1.1 | 0.301 | | | |
| Height at 14 months | Growth at 8 months | $F_{1, 220.2} = 421.9$ | **<0.001** | 0.67 | 0.74 | lmer(height4 ~ grow1 + region + (1 \| cohort) + (1 \| rep2)) |
| | Region | $F_{1, 18.6} = 4.5$ | **0.047** | | | |
| | Cohort(Region) | $\chi^2$ (1) = 21.4 | **<0.001** | | | |
| | Block2 | $\chi^2$ (1) = 1.4 | 0.241 | | | |
| Height at 20 months | Growth at 8 months | $F_{1, 226.4} = 307.2$ | **<0.001** | 0.58 | 0.70 | lmer(height5 ~ grow1 + region + (1 \| cohort) + (1 \| rep2)) |
| | Region | $F_{1, 18.4} = 5.6$ | **0.029** | | | |
| | Cohort(Region) | $\chi^2$ (1) = 37.5 | **<0.001** | | | |
| | Block2 | $\chi^2$ (1) = 1.2 | 0.266 | | | |
| Height at 24 months | Growth at 8 months | $F_{1, 222.6} = 237.5$ | **<0.001** | 0.51 | 0.65 | lmer(height6 ~ grow1 + region + (1 \| cohort) + (1 \| rep2)) |
| | Region | $F_{1, 18.6} = 7.5$ | **0.013** | | | |
| | Cohort(Region) | $\chi^2$ (1) = 39.0 | **<0.001** | | | |
| | Block2 | $\chi^2$ (1) = 0.5 | 0.483 | | | |
| Total growth | Growth at 8 months | $F_{1, 1089.2} = 2122.4$ | **<0.001** | 0.83 | 0.87 | lmer(grow ~ grow1 + time*region + (1 \| cohort) + (1 \| rep2)) |
| (repeated measures) | Time | $F_{4, 1130.0} = 910.0$ | **<0.001** | | | |
| | Region | $F_{1, 18.2} = 0.2$ | 0.671 | | | |
| | Time*Region | $F_{4, 1130.0} = 0.7$ | 0.625 | | | |

| | | | | | | |
|---|---|---|---|---|---|---|
| | Cohort(Region) | $\chi^2$ (1) = 146.6 | **<0.001** | | | |
| | Block2 | $\chi^2$ (1) = 50.6 | **<0.001** | | | |
| Total growth at 24 months | Growth at 8 months | $F_{1, 226.5}$ = 452.3 | **<0.001** | 0.69 | 0.77 | lmer(grow6 ~ grow1 + region + (1 \| cohort) + (1 \| rep2)) |
| | Region | $F_{1, 18.6}$ = 0.0 | 0.844 | | | |
| | Cohort(Region) | $\chi^2$ (1) = 28.0 | **<0.001** | | | |
| | Block2 | $\chi^2$ (1) = 3.8 | 0.051 | | | |
| Biomass at 24 months | Growth at 8 months | $F_{1, 227.3}$ = 422.8 | **<0.001** | 0.69 | 0.80 | lmer(biomass ~ grow1 + region + (1 \| cohort) + (1 \| rep2)) |
| | Region | $F_{1, 18.5}$ = 7.1 | **0.015** | | | |
| | Cohort(Region) | $\chi^2$ (1) = 35.9 | **<0.001** | | | |
| | Block2 | $\chi^2$ (1) = 19.2 | **<0.001** | | | |
| Biomass:Height at 24 months | Growth at 8 months | $F_{1, 219.3}$ = 97.0 | **<0.001** | 0.47 | 0.61 | lmer(bio.h ~ grow1 + region + (1 \| cohort) + (1 \| rep2)) |
| | Region | $F_{1, 18.3}$ = 21.5 | **<0.001** | | | |
| | Cohort(Region) | $\chi^2$ (1) = 19.3 | **<0.001** | | | |
| | Block2 | $\chi^2$ (1) = 10.2 | **0.001** | | | |
| Biomass:Growth at 24 months | Growth at 8 months | $F_{1, 220.9}$ = 25.9 | **<0.001** | 0.23 | 0.44 | lmer(bio.gr ~ grow1 + region + (1 \| cohort) + (1 \| rep2)) |
| | Region | $F_{1, 18.1}$ = 9.6 | **0.006** | | | |
| | Cohort(Region) | $\chi^2$ (1) = 20.5 | **<0.001** | | | |
| | Block2 | $\chi^2$ (1) = 12.0 | **<0.001** | | | |
| Leaf area ($cm^2$) | Growth at 8 months | $F_{1, 215.7}$ = 0.0 | 0.931 | 0.00 | 0.29 | lmer(LA ~ grow1 + region + (1 \| cohort) + (1 \| rep2)) |
| | Region | $F_{1, 18.7}$ = 0.0 | 0.844 | | | |
| | Cohort(Region) | $\chi^2$ (1) = 39.8 | **<0.001** | | | |
| | Block2 | $\chi^2$ (1) = 0.1 | 0.711 | | | |
| Leaf dry-matter content (g/g) | Growth at 8 months | $F_{1, 220.3}$ = 8.4 | **0.004** | 0.04 | 0.33 | lmer(LDMC ~ grow1 + region + (1 \| cohort) + (1 \| rep2)) |
| | Region | $F_{1, 18.3}$ = 1.4 | 0.251 | | | |

| | | | | | | |
|---|---|---|---|---|---|---|
| | Cohort(Region) | $\chi^2$ (1) = 25.8 | **<0.001** | | | |
| | Block2 | $\chi^2$ (1) = 11.9 | **<0.001** | | | |
| Specific leaf area ($cm^2/g$) [a] | Growth at 8 months | $F_{1,\ 210.0}$ = 22.0 | **<0.001** | 0.36 | 0.56 | lmer(SLA ~ grow1 + region + (1 \| cohort) + (1 \| rep2)) |
| | Region | $F_{1,\ 19.0}$ = 51.2 | **<0.001** | | | |
| | Cohort(Region) | $\chi^2$ (1) = 20.0 | **<0.001** | | | |
| | Block2 | $\chi^2$ (1) = 22.2 | **<0.001** | | | |
| log(C:N) [a] | Growth at 8 months | $F_{1,\ 179.1}$ = 0.0 | 0.923 | 0.11 | 0.33 | lmer(l.CNr ~ grow1 + region + (1 \| cohort) + (1 \| rep2)) |
| | Region | $F_{1,\ 17.0}$ = 12.9 | **0.002** | | | |
| | Cohort(Region) | $\chi^2$ (1) = 7.4 | **0.006** | | | |
| | Block2 | $\chi^2$ (1) = 16.3 | **<0.001** | | | |
